# Activity-dependent circuitry plasticity via the regulation of the histamine receptor level in the *Drosophila* visual system

**DOI:** 10.1101/2021.11.22.469494

**Authors:** Yiming Bai, Takashi Suzuki

## Abstract

Activity-dependent synaptic plasticity is crucial for responses to the environment. Although the plasticity mechanisms of presynaptic photoreceptor neurons in the *Drosophila* visual system have been well studied, postsynaptic modifications remain elusive. In addition, further studies on the adaption of the visual system to different light experiences at a circuitry scale are required. Using the modified transcriptional reporter of intracellular Ca^2+^ method, we describe a way to visualize circuitry changes according to different light experiences. We found enhanced postsynaptic neuronal activity responses in lamina monopolar neuron L2 after prolonged light treatment. Although L1 also has connections with photoreceptors, there were no enhanced activity responses in L1. We also report in this study that activity-dependent transcriptional downregulation of inhibitory histamine receptors (HRs) occurs in postsynaptic neuron L2, but not in L1, during continuous light conditions. We expressed exogenous HR proteins in L2 neurons and found that it attenuated the enhanced activity response caused by constant light exposure. These findings, together with the fact that histamine is the main inhibitory neurotransmitter released by photoreceptors in the *Drosophila* visual system, confirmed our hypothesis that the activity-dependent transcriptional downregulation of HRs is responsible for the constant light exposure-induced circuitry response changes in L2. The results successfully demonstrated the selective circuit change after synaptic remodeling evoked by long-term activation and provided *in vivo* evidence of circuitry plasticity upon long-term environmental stimulation.

## Introduction

The ability of animals to adapt to their environment by modifying the structure, connections, or molecular process in their brains is known as “plasticity.” Plasticity allows animals to develop mature brains, recover from injury, and respond to various stimuli and experiences in the environment. Activity-dependent synaptic plasticity, a component of activity-dependent neuroplasticity in the nervous system, is believed to be crucial for responses to the environment and adaptive behaviors. Activity-dependent synaptic plasticity is adopted through the adjustment of the connection strength or information flow according to environmental stimulation.

Many studies have been conducted to investigate the activity-dependent synaptic plasticity that occurs between one neuron and another. For example, in the mammalian central nervous system, activity-dependent synaptic plasticity regarding long-term potentiation (LTP) or long-term depression (LTD) has been well studied (Song and L.Huganir, 2002; Ho et al., 2011). Numerous pre- and postsynaptic mechanisms of plasticity have been studied at excitatory chemical synapses in the rodent hippocampus (Ho et al., 2011; Buonarati et al., 2019), including reorganization of synaptic components, regulation of neurotransmitter release, and postsynaptic regulation of receptors (Lisman et al., 2002; Okamoto et al., 2004; Lee et al., 2009). However, further study is required to understand how synaptic plasticity affects circuitry changes. When a circuit receives an input activity, direct synaptic changes that have subsequent consequences in the circuit occur. Because one type of presynaptic neuron usually has connections with various types of postsynaptic neurons, it remains a big question whether synaptic plasticity happens among all connections or whether only some specific subsets of postsynaptic neurons experience changes in the connections with the presynaptic neuron and modify their activity responses according to the input activity.

Our research aimed to visualize the circuitry changes after synaptic modification evoked by long-term activation and to find *in vivo* evidence of circuitry plasticity after long-term environment stimulation. To accomplish this, we focused on the visual system of the model animal *Drosophila melanogaster.* The *Drosophila* visual system, either in the developing or adult stage, is a powerful model to study the activity-dependent synaptic plasticity evoked by external stimulation from the environment (Yuan et al., 2011; Sugie et al., 2018). Many sophisticated genetic tools and experimental methods have already been developed and applied in the *Drosophila* visual system. Moreover, the small-sized brains, well studied genome and anatomical structures, and advanced imaging techniques make the *Drosophila* visual system an advantageous model for elucidating neural circuit changes.

The adult *Drosophila* visual system comprises the retina and optic lobe. The optic lobe consists of the lamina, medulla, lobula, and lobula plate. The retina consists of approximately 750 ommatidia (small eyes), and each ommatidium has eight types of photoreceptor neuron (R1 to R8). R1 to R6 are responsible for light sensing and motion detection, R7 expresses UV-sensitive opsins, and R8 expresses blue- and green-sensitive opsins (Alejevski et al., 2019). R8 is also believed to be involved in the color-sensing function. The R1–6 photoreceptors project to the lamina, and the R7 and R8 photoreceptors extend to the M6 and M3 layers of the medulla, respectively. Photoreceptors in *Drosophila* mainly release the inhibitory neurotransmitter histamine to their postsynaptic neurons (Alejevski et al., 2019). In the mammalian brain, histamine receptors (HRs) are mostly G protein-coupled receptors (Hill et al., 1997). In *Drosophila,* however, HRs in neurons postsynaptic to photoreceptors is histamine-gated chloride channels. Receiving histamine can cause hyperpolarization in the neuron, thereby decreasing activity.

Previous research has shown that in the adult *Drosophila* visual system, under continuous light conditions (LL), photoreceptor neurons, especially R8, reorganize synaptic components at the active zone (AZ) through microtubule destabilization (Sugie et al., 2015). The activities of both pre- and postsynaptic neurons are involved in synaptic component reorganization. This type of synaptic component reorganization can be reversed when flies are returned to continuous dark conditions (DD) or normal light–dark conditions (LD) for 72 h. However, the subsequent influence of long-term light stimulation on the circuit remains poorly understood. Moreover, although the presynaptic process during synaptic plasticity has been intensely studied, we know significantly less regarding the postsynaptic process in the adult *Drosophila* visual system.

In this research, we applied a transcriptional reporter of intracellular calcium (TRIC) (Gao et al., 2015) to the neurons postsynaptic to photoreceptors to monitor circuitry response and found selective response changes to subsequent environmental stimulation after constant light exposure. Surprisingly, only a subset of postsynaptic neurons (such as lamina monopolar neuron L2) tended to show a drastic activity response to 1-day light stimulation after constant light exposure. Although lamina monopolar neuron L1 also has connections with photoreceptors, it showed no enhanced activity response.

Regarding the underlying mechanism of the selective response changes in the circuit, we hypothesized that the expression levels of HRs in postsynaptic neurons may change because histamine is the main inhibitory neurotransmitter released by photoreceptors in the *Drosophila* visual system. We conducted a number of genetic experiments and found that in the adult *Drosophila* visual system, after constant light exposure, transcriptional downregulation of HRs occurs in postsynaptic lamina neuron L2. The transcriptional downregulation of HRs depends on the photoreceptor and postsynaptic neuronal activity, and the process involves CaMKII and CREB-B. Our research elucidated the activity-dependent synaptic plasticity in the postsynaptic terminal during constant light exposure in the *Drosophila* visual system. The transcriptional downregulation of HRs results in an enhanced postsynaptic L2 neuronal response to subsequent light stimulation. Because HR downregulation happens in L2 but not in L1 during constant light exposure, the circuit shows selective activity response changes. The results successfully demonstrated the circuit change after synaptic remodeling evoked by long-term activation and revealed *in vivo* evidence of the circuitry plasticity upon long-term environmental stimulation. The findings in this research deepen the understanding of the consequences of activity-dependent synaptic plasticity to the circuit and provide new insights into the activity-dependent synaptic plasticity mechanism in the neural circuit.

## Material and Methods

### Fly husbandry

Flies were raised in vials containing a 2 cm cornmeal, yeast-based food layer. The ingredients of fly food and amounts were as follows: water (1L), soybean (15g), agar (10g), cornmeal (100g), yeast (30g), malt (30g) and glucose (50g). The mixture was heated at 90°C for 1 hour. After that, 5 mL ethanol, 2 mL propionic acid, and 2.5 g Nipagin were added, and then the mixture was dispensed into vials and bottles. The vials were placed in plastic tray vial containers and stored in different incubators depending on purposes. For expansion, fly lines were kept in 25°C and 60% humidity incubators with the 12-hour light, 12-hour dark cycle, also known as LD. The fly stocks were kept at 18°C dark room. The temperature and humidity of the incubators were kept at a constant value.

### Light stimulation

The environment change was stimulated as light conditions change. Normally, the flies were reared in 12-hour light, 12-hour dark cycle. For generating the constant condition, we reared the flies in,

1. 24-hour dark (continuous dark/DD) in 25°C or 18°C incubator
2. 24-hour light (continuous light/LL) in 25°C or 18°C incubator [light intensity 4000 lux]

### TRIC assay and GAL80^ts^

Flies carrying TRIC components (LexAop2-mCD8GFP, UAS-mCD8RFP; nSyb-nlsLexADBDo; UAS-p65ADCaM) were crossed with GAL4 lines. On day 3, adult flies were transferred, and vials were covered by foil paper, placed at 25°C for 8 days until eclosion. On day 10, flies were collected and divided into several groups according to experimental requirements.

GAL80^ts^, a temperature-sensitive version of the GAL80 protein, functions as a repressor of GAL4, and temperature-sensitive regulation of GAL4 activity can be realized by conformation change of GAL80^ts^ in different temperatures. GAL80^ts^ is used to restrict reporter availability of the TRIC system until eclosion (McGuire et al., 2003). Flies carrying TRIC components (LexAop2-mCD8GFP, UAS-mCD8RFP; nSyb-nlsLexADBDo; UAS-p65ADCaM) were crossed with GAL4 lines which contained GAL80^ts^ (w; GAL80^ts^/CyO; X-GAL4/TM6B). On day 3, adult flies were transferred, and vials were covered by foil paper, placed at 25°C for 2 days. On day 5, the vials were moved to an 18°C incubator. On day 15, flies were collected and divided into several groups according to experimental requirements (Fig. 1B).

**Fig.1.**
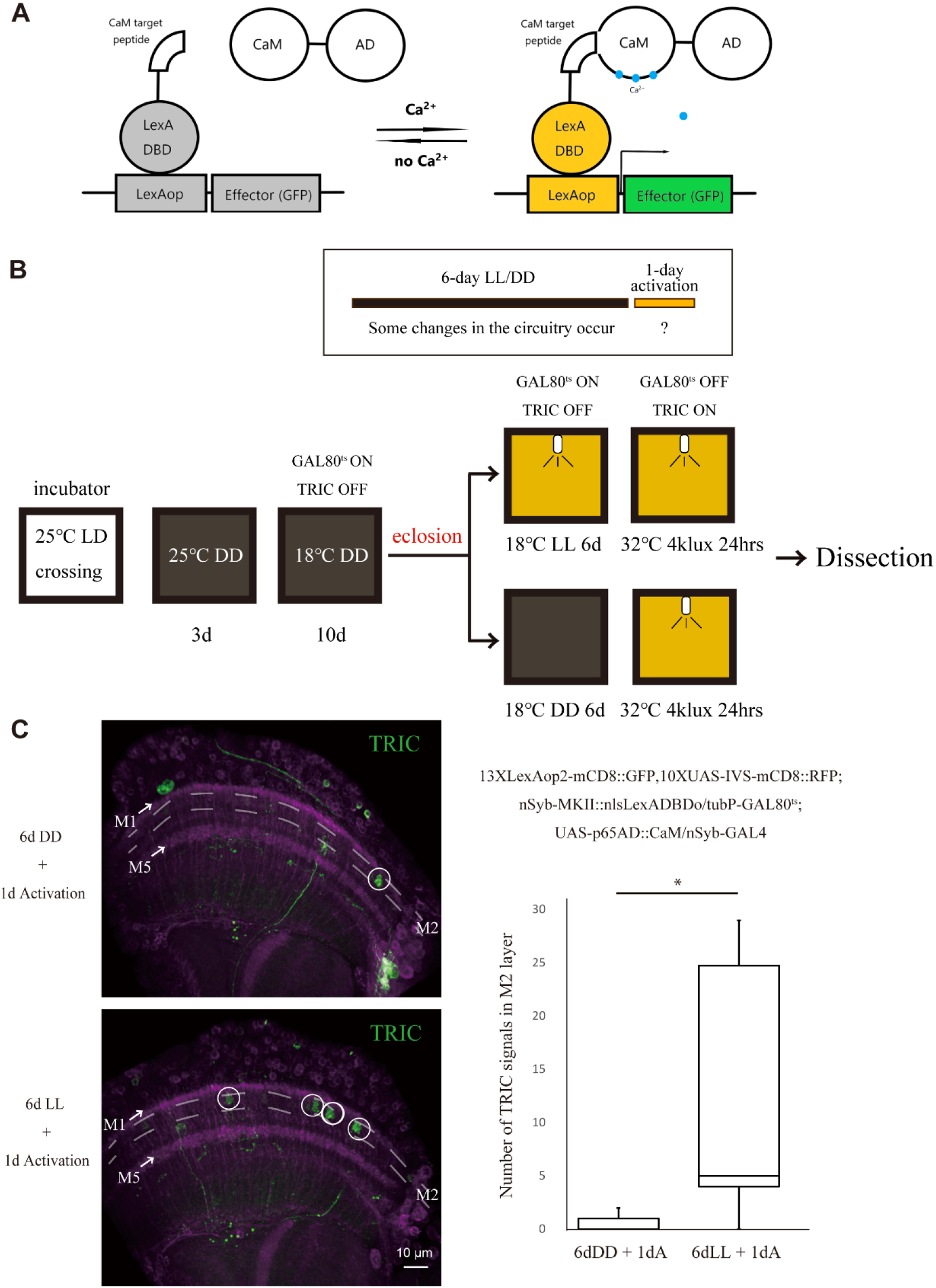
Enhanced activity responses in a subset of postsynaptic neurons after 6-day LL treatment in the TRIC experiment. (A) Scheme of TRIC. TRIC (Transcriptional Reporter of Intracellular Ca^2+^), is a new tool to monitor changes in calcium levels over long periods of time with a split binary expression system. When calcium influx occurs, calmodulin interacts with its target peptides, and allows subsequent expression of reporter genes through the LexA/LexAop system. (B) Experiment design for TRIC assay. After 2-day crossing in 25°C, on Day 3, adult flies were taken away, and vials were covered by foil paper, placed in 25°C for 2 days. On Day 5, the vials were moved to an 18°C incubator. On Day 15, flies were collected and divided into two groups to be treated by LL or DD for 6 days under 18°C after eclosion. 1-day light condition under 32°C was applied on Day 7, and after that fly brains were dissected to monitor the circuitry response towards this 1-day photoreceptor activation. (C) Pan-neuronal TRIC assay. Pan-neuronal TRIC signals in visual circuit for 1-day activation after 6-day LL or DD. Some neurons with oval structure (white circle) in the M2 layer (marked by broken lines) of medulla, mostly like lamina monopolar neuron L2, showed significantly more TRIC GFP signals with step size of 0.5 μm in a sample cross section size of 15μm after 1-day activation subsequent to 6-day LL treatment (12.5 neurons with oval structure in medulla in average) than the ones treated with 6-day DD (0.44 neurons with oval structure in medulla in average). There was no obvious TRIC signal in M1 or M5 (arrow). Scale bar, 10 μm. Student’s t-test, LL: n=10; DD: n=9. p=0.0102.

### OLLAS staining assay

Flies carrying Ort(FRT.SA.Stop)OLLAS;UAS-FLP (Chen et al., 2014) were crossed with L2-specific GAL4, along with UAS-myr-RFP or not according to experimental requirements. In some experiments, flies with recombined R27G05FLP (pan-lamina FLP) and Ort(FRT.SA.Stop)OLLAS on the second chromosome and other components on the third chromosome were crossed with UAS-myr-RFP;L2-specific GAL4 or L2-specifc LexA, LexAop-mCherry. Anti-OLLAS antibody (Funakoshi Anti-OLLAS 0.5 ml NBP1-06713) was used at the concentration of 1:200 and incubated overnight at 4°C with shaking. During the second staining, Alexa Fluor 488 goat anti-rat IgG(H+L) (Life Technologies) were added (1:400) and incubated at room temperature for 2 hours with shaking. After confocal imaging, measurements were done on the IMARIS2 measurement pro track (Carl Zeiss).

### Fly brain dissection and Immunohistochemistry

Fly brains were dissected in 0.1% PBT which was composed of phosphate buffer saline (PBS) and 0.1% Triton X-100 and fixed in fixation solution (4% paraformaldehyde in 0.1% PBT) for 1 hour. After fixation, brains were washed with 0.1% PBT three times and incubated in PBT for 1 hour at room temperature. Brains were then incubated in primary antibody overnight at 4°C. Following first staining with primary antibody, brains were washed with PBT three times and then incubated in secondary antibody for 1 or 2 hours at room temperature. Following second staining with secondary antibody, brains were washed with PBS two times and mounted in Vectashield (Vector Lab), ready for confocal imaging.

If the experiment does not require immunostaining, after fixation, brains were washed with 0.1% PBT three times and incubated in PBT for 1 hour at room temperature. After that, brains were directly washed with PBS twice and mounted in Vectashield, ready for confocal imaging.

Antibodies used in this research were as follows: mAb24B10 (DSHB, 1:50), anti-OLLAS (Funakoshi, 1:200) and goat anti-rat IgG(H+L) Alexa Flour 488 conjugate (Life Technologies, 1:400).

### Confocal imaging and image quantification

Dissection was carried out using Leica light microscope. All confocal images were captured using Nikon Eclipse Ni microscope and Nikon C2plus Confocal microscope. Scanning was done with a step size of 0.5-1 μm in a sample cross-section size of 15-30 μm. The 3D images are acquired using NIS-Element AR (Nikon). After confocal imaging, quantifications were done using the IMARIS2 measurement pro track (Carl Zeiss).

For histamine receptor intensity quantification, the surfaces of L2 neurons were defined by IMARIS2 via L2-specific RFP or mCherry fluorescence signal, and the average staining green fluorescence signal intensity in L2 was calculated in a weighted average manner based on the volumes of different defined regions. The average intensity of the designated area in lobula was subtracted from the original data to eliminate the staining background.

### Real time PCR

RNA was extracted from 10 flies for each experiment condition and the process followed the protocol of Promega ReliaPrepTM RNA Tissue Miniprep System kit (Promega, Z6110). Extracted RNA was stored in a −70°C refrigerator. ReverTra Ace^®^ qPCR RT Master Mix with gDNA Remover (TOYOBO, FSQ-301) was used to synthesize cDNA from 0.5 μg extracted RNA for real-time PCR.

Real time PCR reaction mixtures were prepared using THUNDERBIRD^®^ SYBR^®^ qPCR Mix (TOYOBO, QPS-201) and the process followed the protocol. Real time PCR was done via Real-time PCR equipment TP970 (Takara) and analyzed by Thermal Cycler Dice^®^ Real Time System (Takara).

Primers were designed to target OLLAS-GS-Ort, which indicated the mRNA level of OLLAS-tagged histamine receptor protein Ort. Three independent biological replicates and several technical replicates for each biological replicates were measured for LL or DD conditions. Housekeeping gene *Rpl32* was used as a reference, and primers were referred from FlyPrimerBank. Real time PCR primers were as follows: OLLAS-GS-Ort forward primer (CCTGATGGGCAAGGGTGG) and reverse primer (CTTCGGCGGTCTCATCTTGT); Housekeeping gene *Rpl32* forward primer (CGGATCGATATGCTAAGCTGT) and reverse primer (CGACGCACTCTGTTGTCG).

The calculation of threshold cycle (Ct) of qPCR followed 2nd Derivative Maximum method. The statistical analysis was applied on 2^-ΔΔCt^.

### Statistical Analysis

Statistical analysis between two groups was performed using a two-tailed Student’s t-test. Statistical analysis among more than two groups was performed using non-parametric Kruskall-Wallis H test (Shapiro-Wilk test for the normality and Levene’s Equality of Variances have been applied), followed by post hoc pairwise Mann-Whitney U tests. All *p* values were corrected according to the Holm-Bonferroni method to control for the false discovery rate within multiple comparisons. Significance is shown by asterisks in figures as follows: \**p*<0.05, \*\**p*<0.01, \*\*\**p*<0.001.

### Fly stocks (Supplementary Materials)

Fly stocks were as follows:

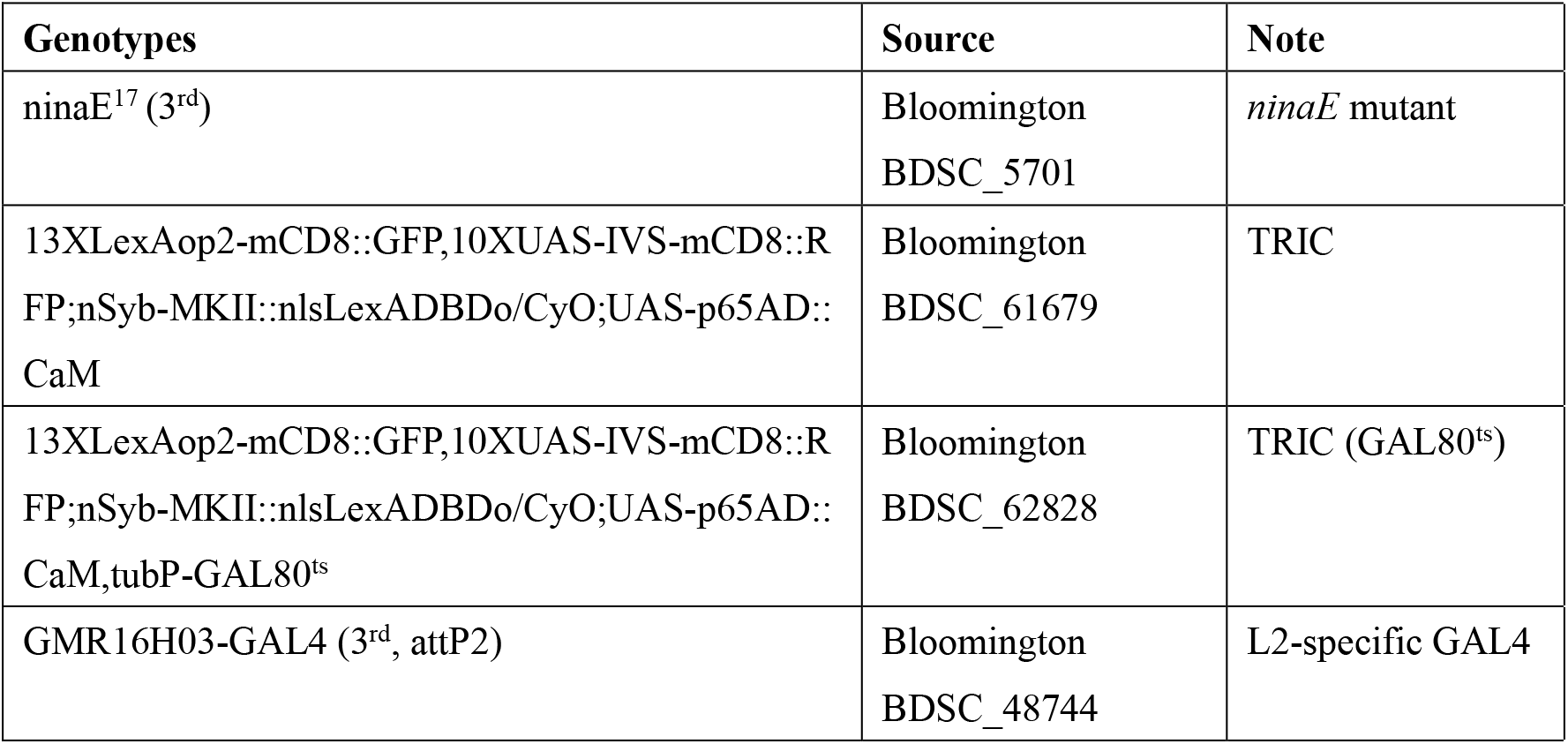
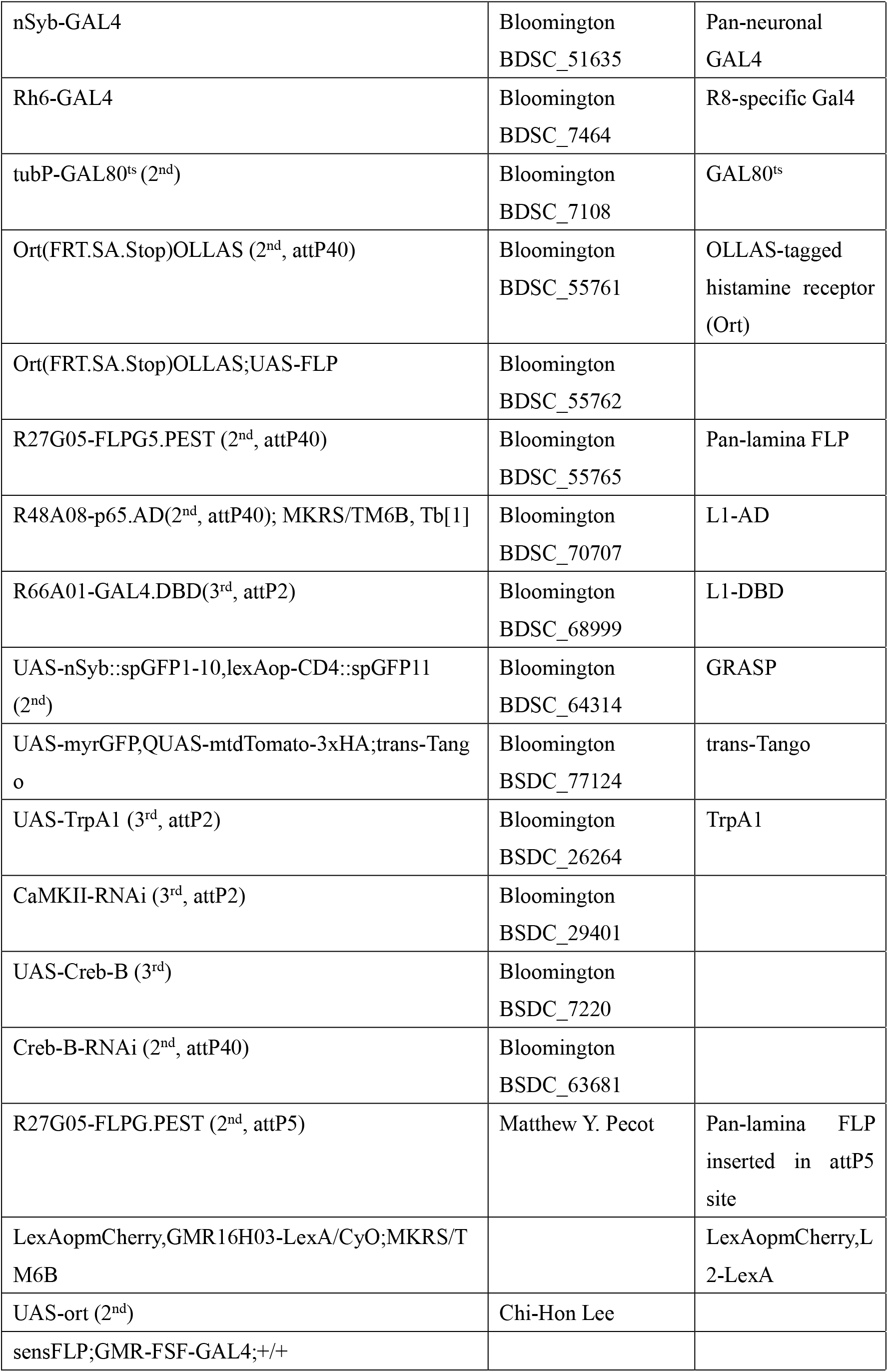

### Genotypes (Supplementary Materials)

The genotypes used in figures were as follows:

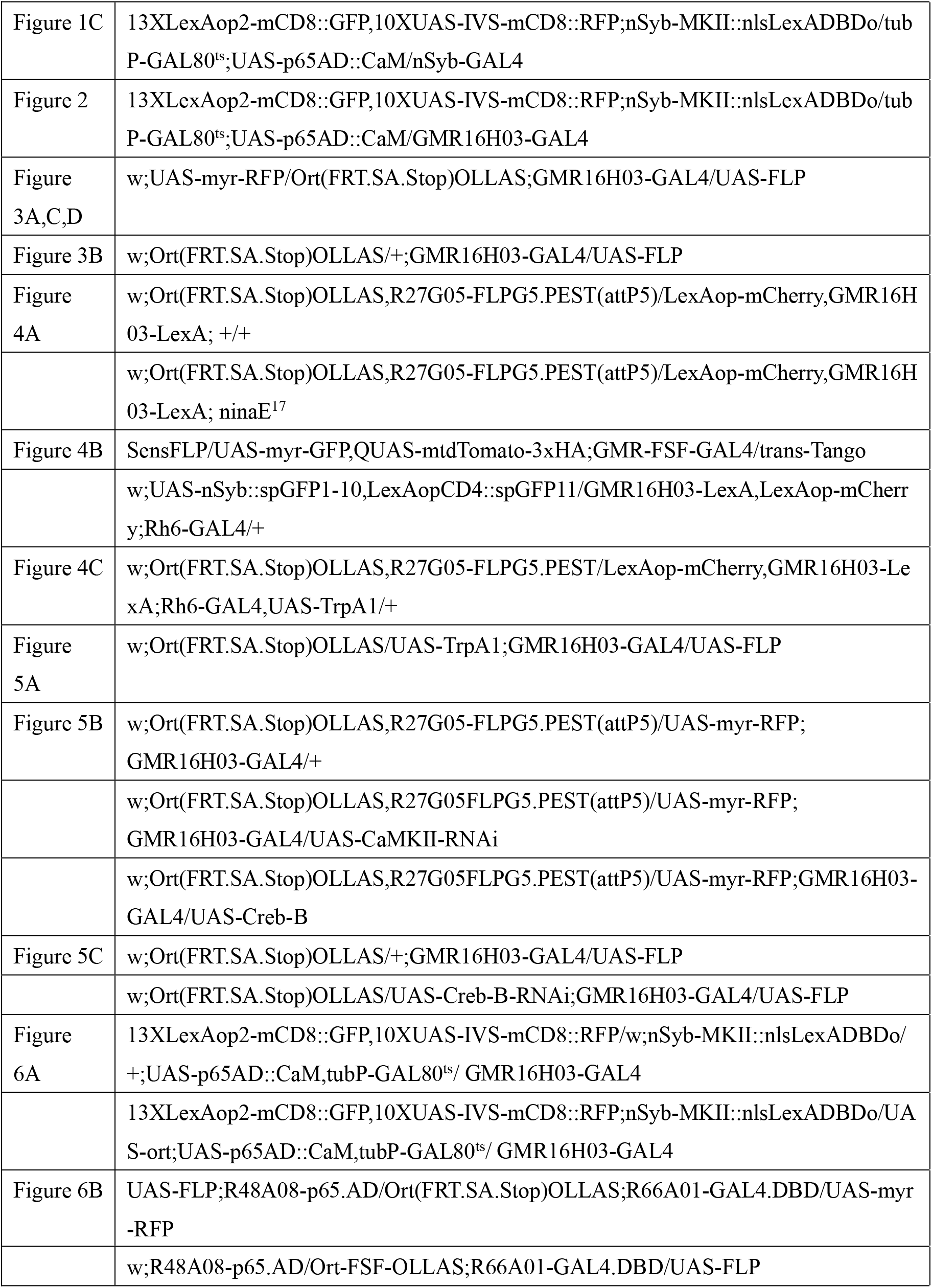

## Results

### Specific postsynaptic neurons in the medulla show enhanced activity responses toward photoreceptor activation after constant light exposure

In a previous study, it was shown that the number of synapses reduced in photoreceptor R8 of *Drosophila* after a 3-day LL condition (Sugie et al., 2015). This reduction mainly occurred in the M2 layer, the second layer of the medulla (Sugie et al., 2015). The finding indicated that presynaptic components can be adjusted to adapt to different light environments. However, it remains unknown what plasticity changes happen in postsynaptic neurons and whether constant light exposure affects the circuitry responses toward subsequent photoreceptor activation. Thus, after 6-day LL or DD treatment, we activated photoreceptors on day 7 and examined the activity response of the visual system at that time.

To measure the activity response of the related circuit, we used the TRIC method, allowing green fluorescent protein (GFP) expression in neurons where calcium influx occurred, with a binary system (Gao et al., 2015). When calcium influx occurs, calmodulin interacts with its target peptides and allows subsequent expression of reporter genes such as GFP (Fig. 1A). To measure only the calcium influx on day 7 and avoid saturation of TRIC signals during the development stage, we restricted TRIC reporter availability until day 7 via the temperature-sensitive GAL80 repressor (GAL80^ts^). GAL80^ts^ functions as an on–off switch for the GAL4-UAS system because it functions as a repressor of GAL4 at 18°C and does not repress GAL4 at temperatures over 29°C (McGuire et al., 2003). Flies were raised according to a modified protocol to avoid a heavy background during the development stage and were subjected to continuous light (LL) or continuous dark conditions (DD) for 6 days at 18°C after eclosion. Then, we activated the photoreceptors via a 1-day continuous light condition at 32°C on day 7 to test the circuitry activity response to this 1-day photoreceptor activation (Fig. 1B). We expressed the TRIC system pan-neuronally and found that some neurons with oval axon terminals in the medulla, mostly likely lamina monopolar neuron L2, showed significantly higher levels of TRIC GFP signals evoked by 1-day light activation after the 6-day LL treatment compared to the ones treated with the 6-day DD condition. After 6 days of LL treatment and 1 day of activation, an average of 12.5 neurons with oval axon terminals in the medulla exhibited TRIC signals within a 15-μm range of depth, and few could be observed after 6 days of DD treatment and 1 day of activation (Fig. 1C). However, although lamina monopolar neuron L1 is postsynaptic to photoreceptors (Takemura et al., 2013), activity responses were not enhanced in L1-like neurons upon 1-day activation after 6-day LL treatment, judged by no notable TRIC signal change in the M1 or M5 layer (Fig. 1C). This result indicates that only a specific subset of postsynaptic neurons in the medulla displays enhanced activity responses after long-term light treatment.

### Lamina monopolar neuron L2 responds more drastically to photoreceptor activation after prolonged light exposure

To further understand the selective change in activity response, we concentrated on L2 neurons and expressed the TRIC system specifically in L2 (GMR16H03-GAL4). Our results showed that there were significantly more L2 neurons exhibiting bright TRIC signals after 1-day photoreceptor activation in flies treated with the 6-day LL condition (an average of 7.8 neurons within a 15-μm range of depth) than in the ones treated with the 6-day DD condition (an average of 1.6 neurons within a 15-μm range of depth) (Fig. 2A). Moreover, these signals were proven not to be derived from incomplete GAL80^ts^ repression because we observed significantly fewer TRIC positive signals when flies were dissected directly without 1-day photoreceptor activation (Fig. 2A). To exclude the possibility that the TRIC signals in the activation stage (day 7) were derived from calcium accumulation during the 6-day LL condition, we also conducted extra TRIC assays in which flies were treated by the condition of 6-day LL and then 1-day DD at 32°C. The results showed that there were significantly fewer TRIC signals in L2 when flies had been treated with 6-day LL and then 1-day 32°C DD compared to when flies had been treated with 6-day LL and then 1-day 32°C LL (Fig. 2B). This result indicated that the TRIC signals in the activation stage (day 7) represented the activity response of L2 toward 1-day light stimulation on day 7 but not the calcium accumulation during 6 days of light exposure.

**Fig.2.**
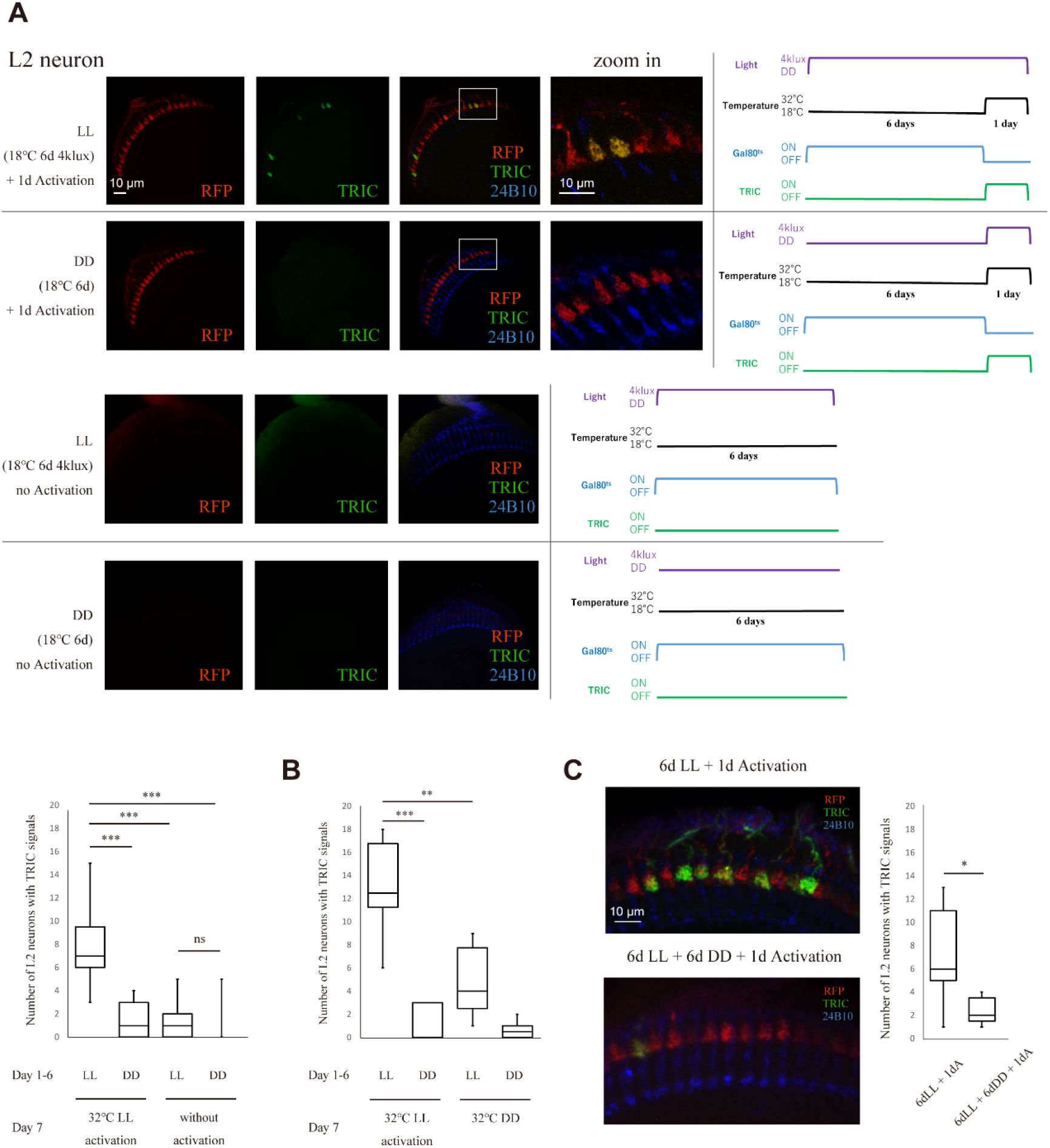
TRIC assay in lamina monopolar neuron L2. (A) TRIC assay specifically performed in L2 neurons of flies (13XLexAop2-mCD8::GFP,10XUAS-IVS-mCD8::RFP; nSyb-MKII::nlsLexADBDo/tubP-GAL80^ts^; UAS-p65AD::CaM/GMR16H03-GAL4). It showed TRIC GFP signals in L2 after 1-day activation subsequent to 6-day LL or DD. Images were taken using NIS-Element AR (Nikon). RFP was used to marked L2, and 24B10 was used to stain photoreceptors. The status of light temperature, Gal80^ts^ and TRIC expression under different conditions are shown beside the confocal images. There were significantly more L2 neurons in flies treated with 6-day LL condition with step size of 0.5 μm in a sample cross section size of 15 μm showing bright TRIC signals after activation than the ones treated with 6-day DD condition. Scale bar, 10 μm. Squared areas were zoomed in to show differences. Statistical tests between groups: p<0.00001, Kruskal-Wallis test; and ***p<0.001, post hoc Mann-Whitney U test. (B) TRIC assays in which flies were treated by conditions as follows: 1) 6-day LL and 1-day 32 °C LL, n=6; 2) 6-day DD and 1-day 32 °C LL, n=5; 3) 6-day LL and 1-day 32 °C DD, n=6; 4) 6-day DD and 1-day 32 °C DD, n=6. Statistical tests between groups: p<0.001, Kruskal-Wallis test; and ***p<0.001, **p<0.01, post hoc Mann-Whitney U test. (C) Reversibility of circuit response change. Response difference caused by continuous light treatment in L2 is reversible. Scale bar, 10 μm. Student’s t-test, 6-day LL + 1-day activation: n=9, avg=7; 6-day LL + 6-day DD + 1-day activation: n=7, avg=2.43. p=0.0119.

We put the flies that had been previously treated with the 6-day LL condition back in the DD environment for 6 more days and observed that there were significantly fewer TRIC GFP signals after 1-day activation (Fig. 2C). This result indicates that prolonged light exposure affects the L2 neuronal response toward subsequent photoreceptor activation and that the response difference caused by continuous light treatment is reversible.

### The histamine receptor level in the medulla arbors of the L2 neurons is regulated by light conditions

Among all neurons, it seemed that only a specific subset of neurons showed an increased activity response toward photoreceptor activation after prolonged light exposure. Other lamina monopolar neurons, such as L1, did not show an increased TRIC positive signal according to the experiment with pan-neuronal expression of TRIC. Therefore, mechanisms within L1 and L2 must work differently under the LL condition. The presynaptic remodeling-based reduction in the synapse number between L2 and the photoreceptors may account for the L2 activity response change (Sugie et al., 2015; Kerwin et al., 2018) because the decreased number of synapses reduced neurotransmitter release from photoreceptors. However, because presynaptic remodeling with a slight reduction in the number of synapses cannot solely explain the robust circuitry response change in postsynaptic neurons caused by activity-dependent synaptic plasticity, we posited that the L2 activity response change could be derived from the histamine receptor reduction in the postsynaptic neurons. We hypothesized that because histamine is an inhibitory neurotransmitter released from *Drosophila* photoreceptors (Dau et al., 2016) and histamine receptors in *Drosophila* are histamine-gated chloride channels, the reduced number of HRs might trigger more calcium influx in L2.

We used OLLAS (OmpF linker and mouse langerin fusion sequence) as a tag linked with the Ort protein (HR protein in postsynaptic neurons of photoreceptors) to label the HR (Chen et al., 2014). To achieve neuron-type-specific expression of the tagged Ort proteins, an FRT-flanked transcriptional and translational stop cassette with a splice acceptor was inserted upstream of the OLLAS-tagged *ort* gene. This stop cassette can only be removed when it is combined with a neuron-type-specific FLP recombinase, allowing the expression of OLLAS-tagged ort protein (Chen et al., 2014) (Fig. 3A). In this experiment, by expressing FLP specifically in L2 neurons, tagged HRs were observed only in L2 via OLLAS staining. The result showed that the intensity of OLLAS signals in the medulla arbors of L2 was significantly decreased under the 6-day LL condition compared to the 6-day DD condition. However, there was no significant decrease in L2-specific tagged HRs after 1- or 3-day LL exposure, indicating that HR downregulation occurred after at least 3 days (Fig. 3A).

**Fig.3.**
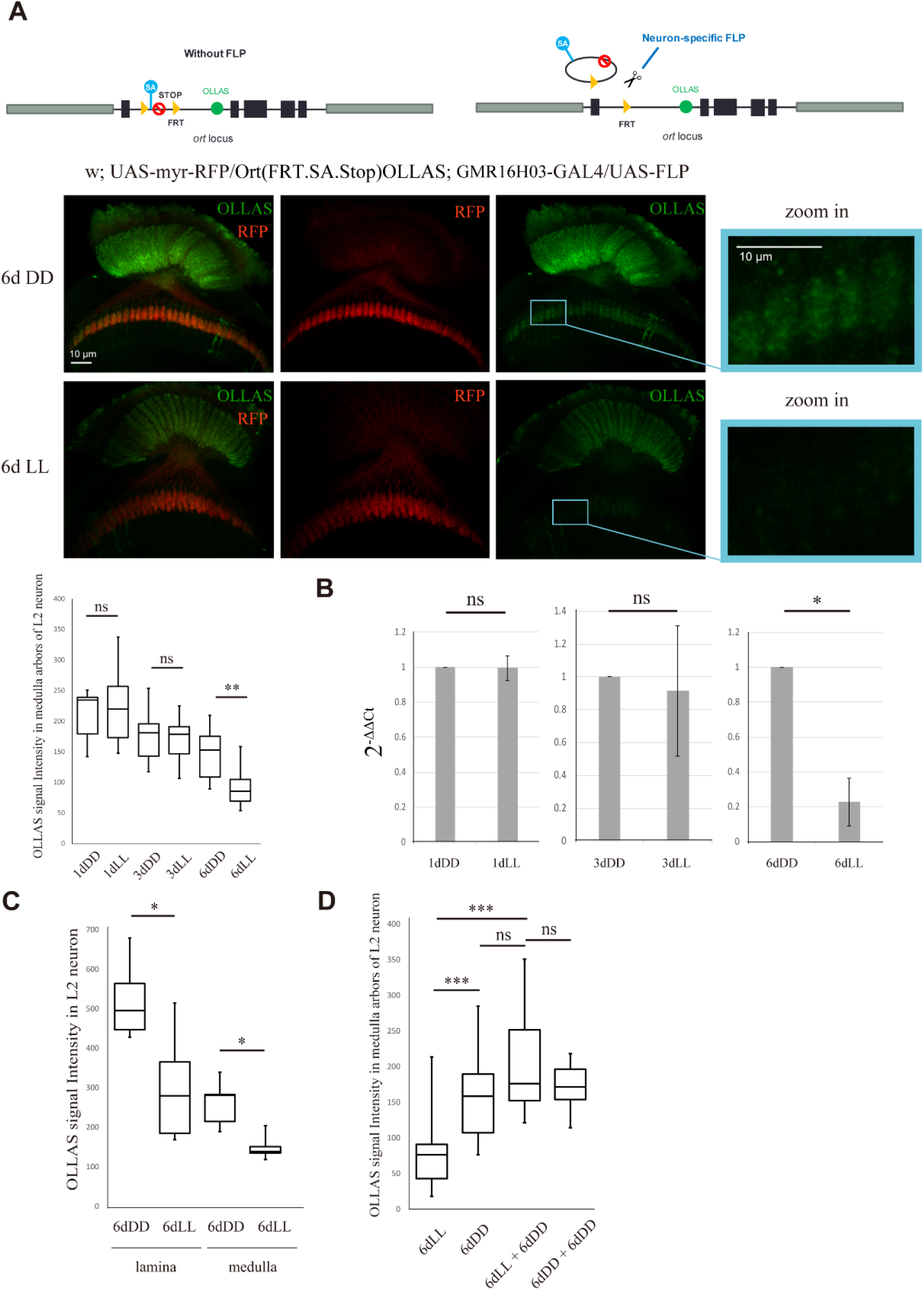
Transcriptional downregulation of histamine receptor in L2. (A) OLLAS (ompF linker and mouse langerin fusion sequence) is used as a tag linked with ORT protein (HR protein in postsynaptic neurons of photoreceptors) to label the HR. To achieve neuron-type-specific expression of tagged Ort proteins, FRT-flanked transcriptional and translational stop cassette with a splice acceptor (SA) was inserted in the upstream of OLLAS-tagged Ort gene. Only when combined with a neuron-type-specific FLP recombinase can the stop cassette be removed, allowing the expression of OLLAS-tagged Ort protein. The illustration is modified from Chen et al., 2014. Immunostaining using an anti-OLLAS antibody (green) of *Drosophila* brains after 6-day LL or DD. Scanning was done with step size of 0.5 μm in a sample cross section size of 15 μm. The intensity of OLLAS signal of L2 in medulla was significantly decreased under 6-day LL condition compared to 6-day DD, while there was no significant decrease after 1-day or 3-day LL. Scale bar, 10 μm. Statistical tests between groups: p=0.00004313, Kruskal-Wallis test; and **p<0.01, post hoc Mann-Whitney U test. (B) Three independent biological replicates (with technical replicates) were measured after LL or DD condition. Housekeeping gene *Rpl32* was used as the reference. The calculation of threshold cycle (Ct) of qPCR followed 2nd Derivative Maximum method. The statistical analysis was applied on 2^-ΔΔCt^. Student’s t-test, 1-day LL: 0.995; 3-day LL: 0.914; 6-day LL: 0.228; p=0.954 (1dDD–1dLL), p=0.848 (3dDD–3dLL), p =0.0298 (6dDD–6dLL). (C) Pattern of HR transcriptional downregulation in L2. OLLAS staining intensity changed after LL/DD in lamina and medulla parts of L2 respectively. Statistical tests between groups: p=0.001433, Kruskal-Wallis test; and *p<0.05, post hoc Mann-Whitney U test. (D) Downregulation of histamine receptor in medulla part of L2 was reversible. Statistical tests between groups: p=0.00001755, Kruskal-Wallis test; and ***p<0.001, post hoc Mann-Whitney U test.

To investigate whether this kind of HR downregulation is transcriptional, reverse transcriptional PCR and real-time PCR were applied to measure the mRNA level of L2-specific tagged HRs. Primers designed to target a fragment of OLLAS-Ort (one primer on OLLAS and the other on Ort) were used to conduct real-time qPCR. Although the transcription of tagged HR in other tissues was prevented by the FRT-STOP-FRT cassette, the tagged HR in which the FRT-STOP-FRT cassette had been removed by L2-specific FLP could be successfully transcribed. Thus, only the mRNA level of tagged HR specifically in L2 could be measured, even though the mRNA was extracted from the entire body. These results showed that the transcription level of L2-specific OLLAS-Ort was significantly decreased after 6-day LL treatment compared with 6-day DD treatment. The mRNA level of the L2-specific OLLAS-Ort protein after 6 days of LL treatment was only 22.8% of that after 6 days of DD treatment (Fig. 3B). This result showed that this kind of HR downregulation is transcriptional. However, after 3- or 1-day LL, the mRNA level of L2-specific OLLAS-Ort showed no significant difference compared with that of 3- or 1-day DD, respectively (Fig. 3B). This result indicates that transcriptional regulation of OLLAS-Ort proteins starts no earlier than 3 days. Prolonged light exposure lasting longer than 3 days is therefore necessary for the regulation of HRs in L2. Moreover, OLLAS staining showed that the GFP intensity in the L2 neurons of flies treated by 6-day LL decreased significantly both in the lamina and medulla (Fig. 3C). This result demonstrates that the HR level is downregulated in both the lamina and the medulla L2 neurons after prolonged light exposure. The decrease in HRs in the medulla could indicate overall HR downregulation in the L2 neurons. Moreover, this type of downregulation was confirmed to be reversible; the decreased HR expression level in L2 due to 6-day LL treatment could be recovered if flies were put back in the DD environment for 6 more days (Fig. 3D).

### R1–6 activities are necessary for histamine receptor loss in the medulla arbors of L2 under continuous light conditions

Among the eight types of photoreceptors, the R1–6 photoreceptors are responsible for light sensing and maintaining connections with L2 in the lamina. The R1–6 photoreceptors are considered the main input sources for L2 neurons. There is a high possibility that the activity of R1–6 accounts for the HR downregulation in L2. Rhodopsin is the protein in the photoreceptors that detects light and vision. In R1–6 photoreceptors, the rhodopsin type is Rh1 (ninaE). In *ninaE* mutants, in which signals from R1–6 are blocked (Chou et al., 1996), even though the HR protein levels in the medulla arbors of L2 in flies treated with 6-day LL were slightly decreased (by 12.8% on average compared to those treated with 6-day DD), the difference was not significant (Fig. 4A). In this experiment, we measured OLLAS-Ort signals via pan-lamina expression of OLLAS-Ort along with specifically expressing mCherry in L2. The results indicated that R1–6 activities are necessary for HR downregulation in the medulla arbors of L2 under continuous light conditions. The slight decrease in the HR level after LL treatment despite the *ninaE* mutation implies that other factors are involved in the downregulation process.

**Fig.4.**
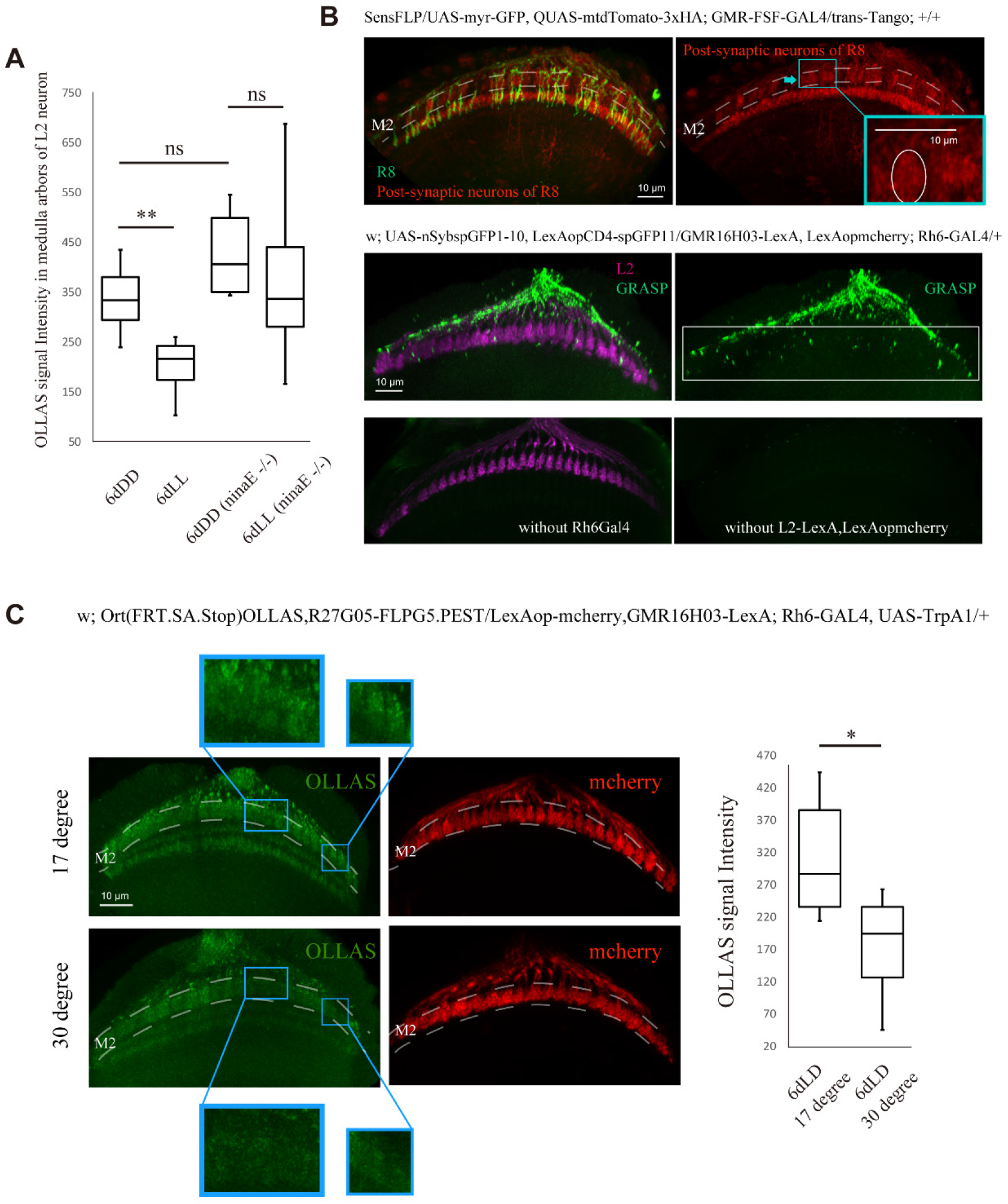
Influence of photoreceptor activities on the HR level in L2. (A) OLLAS intensity has no significant change in L2 neurons of flies with *ninaE* mutant after 6-day LL. *ninaE* mutant: w; Ort(FRT.SA.Stop)OLLAS, R27G05-FLPG5.PEST(attP5) / LexAop-mCherry, GMR16H03-LexA; ninaE^17^. Control: w; Ort(FRT.SA.Stop)OLLAS, R27G05-FLPG5.PEST(attP5) / LexAop-mCherry, GMR16H03-LexA; +/+. Statistical tests between groups: p=0.005843, Kruskal-Wallis test; and **p<0.01, post hoc Mann-Whitney U test. (B) Connection between photoreceptor R8 and L2. Confirmation of the neurons postsynaptic to photoreceptor R8 using trans-Tango revealed L2-like neurons with oval structures (circled area) in the M2 layer (marked by broken lines). Scanning was done with step size of 1 μm in a sample cross section size of 30 μm. Scale bar, 10 μm. Squared areas were zoomed in. Confirmation of the connection between photoreceptor R8 and L2 neurons using activity-dependent GRASP showed green fluorescence signals in medulla (squared area). Scanning was done with step size of 0.5 μm in a sample cross section size of 15 μm. Scale bar, 10 μm. (C) Effect of activating photoreceptor R8 on the HR level in L2 neurons. R8 was artificially activated using TrpA1 and histamine receptor level in some subsets of medullar part of L2 neurons was decreased upon R8 activation in the M2 layer (marked by broken lines). We used mCherry protein to mark L2. Scanning was done with step size of 1μm in a sample cross section size of 20μm. Scale bar, 10 μm. Squared areas were zoomed in to show differences. Student’s t-test, 6-day LD 17°C: n=6, avg=311.76; 6-day LD 30°C: n=6, avg=175.41. p=0.0268.

### Activating photoreceptor R8 is sufficient to induce histamine receptor downregulation in the medulla arbors of L2 under continuous light conditions

According to previous studies, after LL treatment, the synapse number in R8 is decreased mainly in the M2 layer (Sugie et al., 2015). It remains to be confirmed whether R8 photoreceptors have connections with L2 neurons in the M2 layer. Using the trans-Tango approach (Talay et al., 2017), we marked presynaptic R8 photoreceptors with green fluorescence and all neurons postsynaptic to R8 with red fluorescent tandem dimer Tomato (tdTomato) to identify the second-order neurons of R8 and found L2-like neurons with oval axon terminals (Fig. 4B). Using the activity-dependent GFP reconstitution across synaptic partners (GRASP) method, which shows GFP signals at the site of the active synaptic connection between two neurons (Feinberg et al., 2008; Macpherson et al., 2015), we checked the synaptic connections specifically between R8 and L2 and confirmed that R8 photoreceptors have synaptic connections with L2 in the medulla (Fig. 4B). In contrast to our observations, in a previous study, an electron microscopy-assisted connectome reconstruction showed that R8 photoreceptors exhibited no synaptic connections with L2 in the medulla (Takemura et al., 2013). The differences between our findings and those of the previous study could be due to the sparse number of synapses between R8 and L2 because not all R8 photoreceptors seem to maintain connections with the L2 neurons. The reference column and adjacent medullar columns chosen for electron microscopic reconstruction in the previous study may not have included the sparse synaptic connections between R8 and L2 (Takemura et al., 2011; Takemura et al., 2013).

To determine whether the activity of R8 affects the HR level in L2, we artificially activated R8 via TrpA1 (transient receptor potential cation channel A1). TrpA1 is a thermosensitive cation channel that can be activated by warming (Luo et al., 2017). We found that the HR level in the medulla arbors of L2 was decreased upon R8 activation (Fig. 4C). This result indicates that activation of R8 is sufficient to induce HR level downregulation in the medulla arbors of L2 under continuous light conditions.

### The L2 histamine receptor level is regulated by L2 activity via the CaMKII-related pathway

We artificially activated L2 using TrpA1 at 29°C and found severe HR reduction even after 6 days of DD treatment (Fig. 5A). This result indicates that HR downregulation in L2 is activity-dependent. According to previous studies, the *Drosophila* neuromuscular junction and *Drosophila* olfactory system require calmodulin kinase II (CaMKII) for activity-dependent synaptic plasticity (Bai and Suzuki, 2020). Moreover, studies of the rodent hippocampus have demonstrated the involvement of the CaMKII-related pathway and the transcriptional factor cAMP response element-binding (CREB) protein in activity-dependent postsynaptic modification. CREB functions after calcium influx to manipulate the expression of downstream activity-regulated genes (Carlezon et al., 2005). CaMKII is known to inhibit CREB by phosphorylating serine 142 in CREB, which leads to the dissociation of the CREB dimer (Matthews et al., 1994; Wu and McMurray, 2001). In the developing *Drosophila* visual circuit, lipophorin receptors LpR1 and LpR2 in the postsynaptic ventral lateral neurons (LNvs) can be upregulated during constant light exposure (Yin et al., 2018). *Creb-B* is also one of the activity-dependent genes regulated by light exposure (Yuan et al., 2011; Yin et al., 2018).

**Fig.5.**
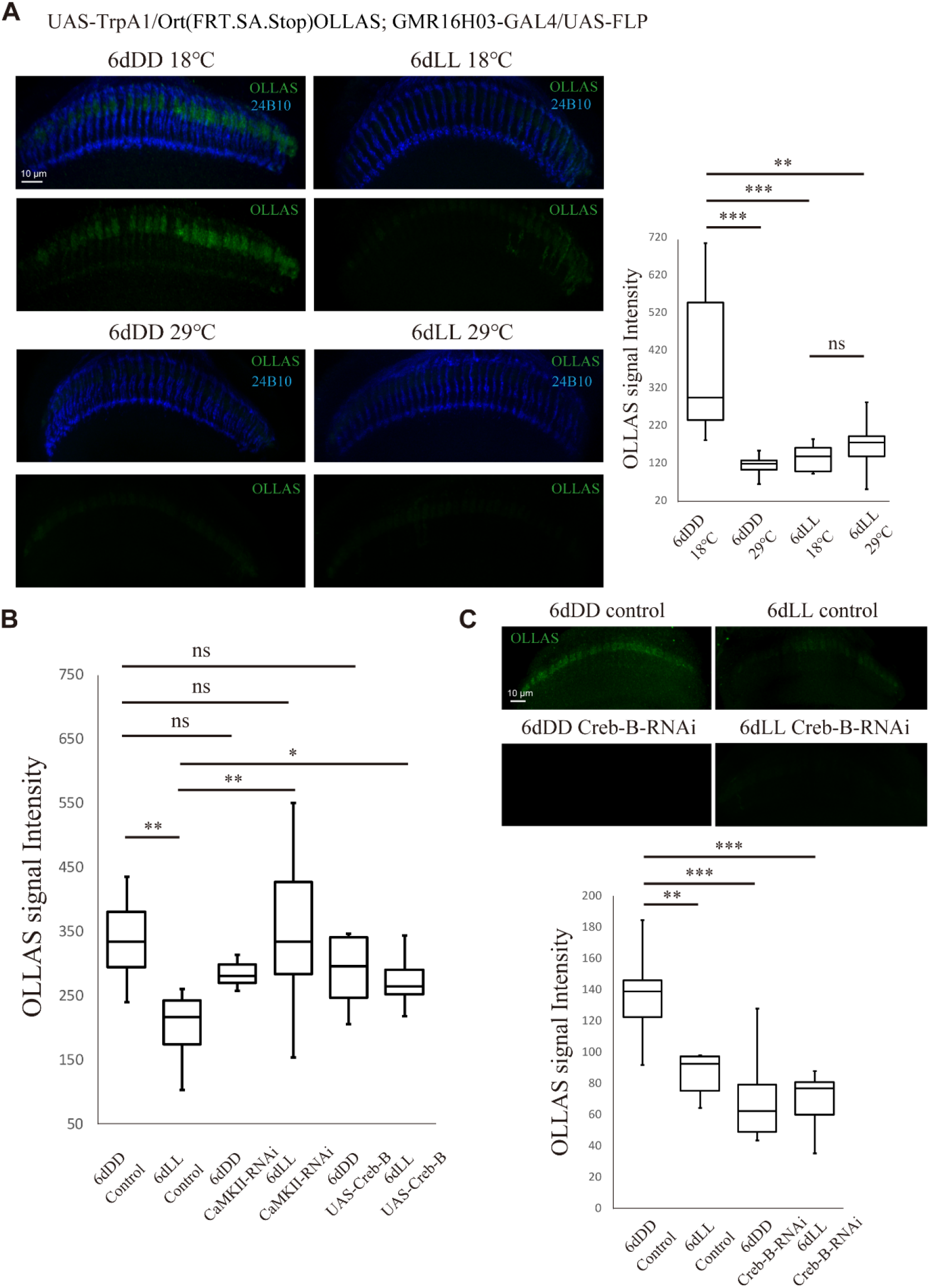
Activity-dependent regulation of histamine receptor level in L2 through CaMKII-related pathway. (A) Artificial activation of L2 with TrpA1 and effect on histamine level. Scanning was done with step size of 1 μm in a sample cross section size of 15 μm. 24B10 was used to stain photoreceptors. Scale bar, 10 μm. Statistical tests between groups: p=0.00002236, Kruskal-Wallis test; and ***p<0.001, **p<0.01, post hoc Mann-Whitney U test. (B) CaMKII-Creb-B pathway. Scanning was done with step size of 0.5 μm in a sample cross section size of 15 μm. Control: w; Ort(FRT.SA.Stop)OLLAS, R27G05-FLPG5.PEST(attP5) / UAS-myr-RFP; GMR16H03-GAL4/+; CaMKII-RNAi: w; Ort(FRT.SA.Stop)OLLAS, R27G05FLPG5.PEST(attP5) / UAS-myr-RFP; GMR16H03-GAL4 / UAS-CaMKII-RNAi; UAS-Creb-B: w; Ort(FRT.SA.Stop)OLLAS, R27G05FLPG5.PEST(attP5) / UAS-myr-RFP; GMR16H03-GAL4 / UAS-Creb-B. Statistical tests between groups: p=0.0151, Kruskal-Wallis test; and **p<0.01, *p<0.05, post hoc Mann-Whitney U test. (C) Knockdown of Creb-B with Creb-B-RNAi largely decrease histamine receptor level in medulla part of L2. Control: w; Ort(FRT.SA.Stop)OLLAS /+; GMR16H03-GAL4 / UAS-FLP; Creb-B-RNAi: w; Ort(FRT.SA.Stop)OLLAS / UAS-Creb-B-RNAi; GMR16H03-GAL4 / UAS-FLP. Scanning was done with step size of 0.5 μm in a sample cross section size of 15 μm. Scale bar, 10 μm. Statistical tests between groups: p=0.000396, Kruskal-Wallis test; and ***p<0.001, **p<0.01, post hoc Mann-Whitney U test.

Considering these findings, we conducted experiments to knockdown CaMKII in L2 with CaMKII-RNAi and found that the HR level was not significantly decreased after 6 days of LL treatment. Overexpression of CREB-B with UAS-Creb-B in L2 also highly suppressed the decrease in HRs in L2 under both continuous light and continuous dark conditions, and the phenotype resembled the expression of CaMKII-RNAi in L2 (Fig. 5B). Knockdown of CREB-B using Creb-B-RNAi enhanced the decrease in HRs in L2 after both 6-day LL and DD conditions (Fig. 5C). Because CaMKII is a negative CREB-B regulator, calcium influx in L2 first stimulates CaMKII, and then, CaMKII suppresses CREB-B, resulting in the downregulation of the expression of HRs in L2. These findings indicate that the CaMKII-related pathway is involved in the activity-dependent transcriptional HR downregulation in L2.

### Exogenous expression of Ort in L2 neurons attenuates the enhanced activity response caused by constant light exposure

Next, using UAS-ort, GAL80^ts^, and the TRIC system, we expressed Ort proteins in L2 neurons on day 7 after 6-day LL treatment and monitored the activity responses of L2 toward 1-day photoreceptor activation simultaneously. The results showed significantly fewer L2 neurons in flies with exogenous expression of Ort in L2 on day 7 after 6-day LL treatment (an average of 2.57 neurons within a 15-μm range of depth) exhibiting TRIC GFP signals after 1 day of photoreceptor activation compared to flies in the control groups treated with 6-day LL and 1-day photoreceptor activation (an average of 7.73 neurons within a 15-μm range of depth) (Fig. 6A). This result demonstrates that exogenous expression of Ort in L2 on day 7 reverted the activity response change caused by prolonged light exposure. Along with the finding that there was transcriptional downregulation of HRs in L2 after 6-day LL treatment, this result supports our hypothesis that the change in L2 activity response to 1-day photoreceptor activation after prolonged light exposure is derived from HR reduction.

**Fig.6.**
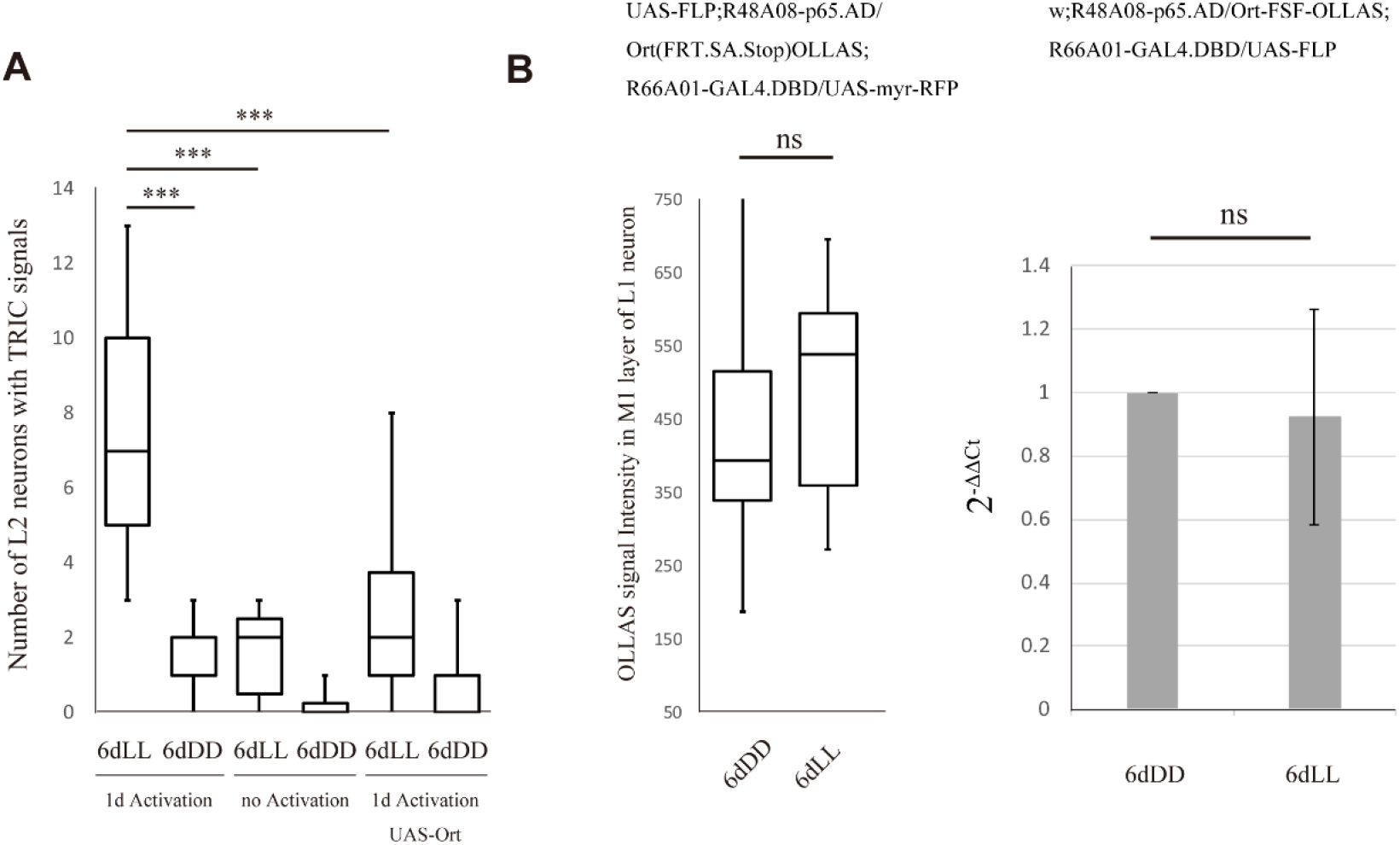
The selective change in activity responses to 1-day photoreceptor activation after prolonged light exposure derived from the HR reduction. (A) TRIC assays in which flies were treated by conditions as follows: 1) 6-day LL and 1-day 32 °C LL, n=11; 2) 6-day DD and 1-day 32 °C LL, n=8; 3) 6-day LL and no activation, n=7; 4) 6-day DD and no activation, n=8; 5) 6-day LL and 1-day 32 °C LL, with UAS-Ort expression, n=14; 6) 6-day DD and 1-day 32 °C LL, with UAS-Ort expression, n=13. Control: 13XLexAop2-mCD8::GFP, 10XUAS-IVS-mCD8::RFP /w; nSyb-MKII::nlsLexADBDo /+; UAS-p65AD::CaM, tubP-GAL80^ts^/ GMR16H03-GAL4. UAS-ort flies: 13XLexAop2-mCD8::GFP, 10XUAS-IVS-mCD8::RFP; nSyb-MKII::nlsLexADBDo /UAS-ort; UAS-p65AD::CaM, tubP-GAL80^ts^/ GMR16H03-GAL4. Statistical tests between groups: p<0.0000001, Kruskal-Wallis test; and ***p<0.001, post hoc Mann-Whitney U test. (B) HR level in L1 neurons under various light conditions. OLLAS intensity has no significant change in L1 neurons of flies after 6-day LL compared to 6-day DD. Student’s t-test, 6-day DD: n=12, avg=496.71; 6-day LL: n=12, avg=491.69. p=0.957. (C) Three independent biological replicates (with technical replicates) were measured for LL or DD condition. Housekeeping gene Rpl32 was used as the reference. The calculation of threshold cycle (Ct) of qPCR followed 2nd Derivative Maximum method. The statistical analysis was applied on 2^-ΔΔCt^. Student’s t-test, 6-day LL: 0.924. p=0.844.

### Histamine receptor downregulation does not occur in L1 upon constant light exposure

The TRIC experiment showed that after prolonged light exposure, only a specific subset of postsynaptic neurons, such as L2 neurons, increases their activity responses upon photoreceptor activation. Although lamina monopolar neuron L1 also has connections with photoreceptors, L1 neurons do not show enhanced activity responses (Fig. 1C). To assess whether the HR downregulation that occurs in L2 after 6-day LL also happens in L1, we checked the HR expression level specifically in L1 and found that there was no significant difference in OLLAS staining intensity in L1 between flies treated with 6-day LL and those treated with 6-day DD (Fig. 6B). Reverse transcriptional PCR and real-time PCR were also applied to examine the mRNA level of OLLAS-tagged HR specifically in L1. The results showed that the transcriptional level of OLLAS-tagged HR was not changed after 6-day LL treatment (Fig. 6B). This result reveals that there is no downregulation of HR in L1 upon prolonged light exposure and explains why activity response changes occurred only in L2 in the TRIC assay.

## Discussion

### The enhanced L2 neuronal activity response is part of the adaption to prolonged light exposure

In this study, we applied the TRIC method to express GFP in neurons postsynaptic to photoreceptors when there was calcium influx to achieve better understanding of the subsequent influence of activity-dependent, environmental stimulation-induced synaptic plasticity on the circuit in the *Drosophila* visual system. The TRIC validation of the physiological response showed that only a subset of postsynaptic neurons, L2 neurons, exhibited a robust activity response to environmental stimulation after chronic light treatment. L1 neurons also have connections with photoreceptors, but their activity responses did not change. This research succeeded in visualizing the circuit change in a selective circuitry after activity-dependent synaptic plasticity evoked by long-term activation.

It has been reported previously that in photoreceptors, ambient light exposure induces the dominant arrestin isoform, Arr2, which is translocated to the rhabdomere loaded with R1–6 and the inactivated photoreceptor response (Satoh et al., 2010). Moreover, recent studies have indicated that continuous light conditions trigger photoreceptor neuronal component reorganization in the AZ through microtubule destabilization and reduction in the number of synapses (Sugie et al., 2015). Such processes allow the *Drosophila* visual system to adapt to prolonged light exposure. Additional to these findings, we propose that the enhanced L2 neuronal activity response is also part of the adaption to prolonged light exposure.

### Postsynaptic modification involving transcriptional regulation of histamine receptors occurs during prolonged light exposure

Activity response changes at synapses can be achieved through postsynaptic modifications, such as regulation of neurotransmitter receptor availability at the postsynaptic terminal (Chater and Goda, 2014; Sugie et al., 2018). Previous studies have shown that the postsynaptic modification during activity-dependent synaptic plasticity probably occurs mainly through the regulation of postsynaptic receptor localization and recycling. For example, in the rodent hippocampus, postsynaptic regulation of AMPA glutamate receptors (AMPARs), which involves the trafficking and recycling of receptors, was found to be crucial for plasticity (Raymond et al., 1993; Song and L.Huganir, 2002; Ho et al., 2011). The exocytosis and endocytosis of AMPARs were found to account for the AMPAR level at the synapse due to LTP or LTD (Buonarati et al., 2019). However, cases of transcriptional postsynaptic receptor regulation remain relatively rare. Moreover, although some previous studies have demonstrated activity-dependent regulation of glutamate receptor localization during the development of *Drosophila* neuromuscular junction (Packard et al., 2003), there are few studies that investigate postsynaptic modification in adult *Drosophila.* In this study, we aimed to discover what happens in the postsynaptic neurons of *Drosophila* photoreceptors during prolonged light exposure and successfully observed a reduction in both HR protein and mRNA levels in the medulla arbors of the L2 neurons under LL conditions. We demonstrated that the HR downregulation in adult *Drosophila* postsynaptic L2 neurons is activity-dependent and occurs on the transcriptional level. Like the postsynaptic modifications in the rodent hippocampus (Carlezon et al., 2005), HR transcriptional downregulation is regulated by CaMKII and CREB-B. These findings supplement the understanding of activity-dependent synaptic plasticity that occurs in the postsynaptic terminal during long-term environmental stimulation.

### Activity-dependent transcriptional downregulation of histamine receptors is responsible for the selective response changes evoked by long-term activation

The results of the *ninaE* mutation experiment showed that R1–6 activities were necessary for HR level regulation in L2 neurons. R1–6 photoreceptors maintain synaptic connections with L2 neurons in the lamina, and L2 neurons mainly receive light information input from R1–6. Therefore, R1–6 photoreceptor activity appears to account for significant HR downregulation in the L2 neurons. Meanwhile, we found synaptic connections between R8 and L2 in the medulla, and activating R8 was sufficient to downregulate L2 HRs, suggesting that the L2 HR level is regulated by multiple factors. R1–6- and R8-related regulatory mechanisms are activity-dependent. In this study, it was also interesting that HR downregulation occurred only in L2 after 6-day LL, but not in L1, which is consistent with the observation that only L2 neurons showed increased activity responses to photoreceptor activation after prolonged light exposure. Exogenous expression of Ort in L2 neurons attenuates the enhanced activity response caused by constant light exposure. These findings, together with the fact that histamine is the main inhibitory neurotransmitter released by photoreceptors in the *Drosophila* visual system, affirm the hypothesis that the activity-dependent transcriptional downregulation of HRs is responsible for the constant light exposure-induced circuitry response change. After the visual circuit of *Drosophila* receives long-term activation, synaptic plasticity involving activity-dependent transcriptional downregulation of HRs occurs specifically in L2 neurons and then changes the activity responses of L2 upon subsequent photoreceptor activation. This implies the special role of L2 neurons in integrating light sensing, motion detection, and color vision, consistent with the central role of L2 as secondary neurons in the medullar visual circuit (Fig. 7).

**Fig.7.**
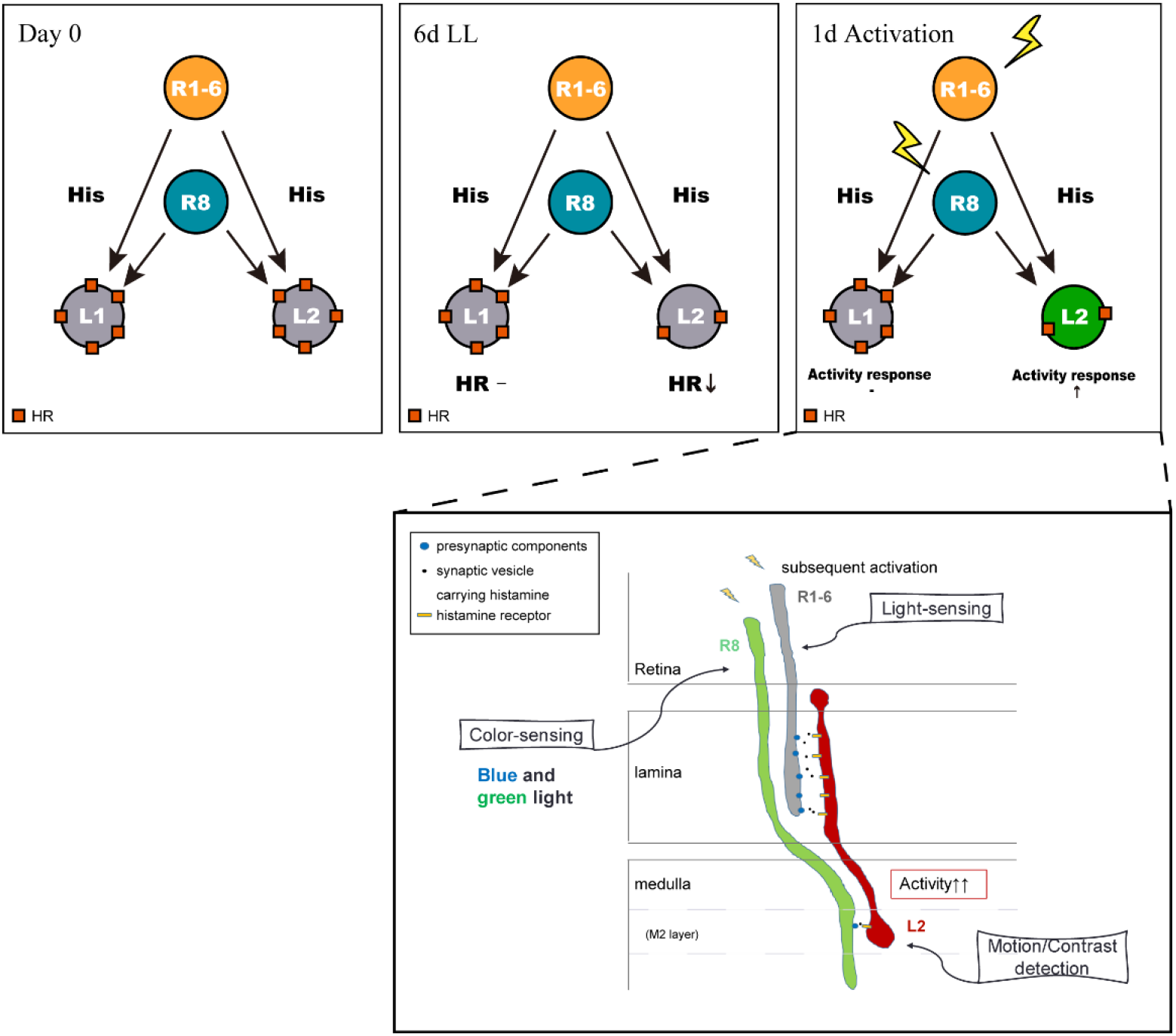
A schematic diagram of the selective circuitry response change after activity-dependent synaptic plasticity evoked by long-term activation in the *Drosophila* visual system. After the visual circuit of Drosophila receives long-term activation (6-day LL treatment), synaptic plasticity involving activity-dependent transcriptional downregulation of HRs occurs specifically in L2 neurons, but not L1, and then enhanced the activity responses of L2 upon subsequent photoreceptor activation on Day 7. This activity-dependent selective circuitry response change implies the process of integrating light-sensing (R1–6), motion detection (L2), and color vision (R8) when flies are faced with prolonged light exposure.

### Time course of activity and histamine receptor regulation in L2

The results of this study showed that HR downregulation in the postsynaptic L2 neurons started no earlier than 3 days after eclosion, which was consistent with the fact that the 3-day LL condition was not sufficient to cause an increased L2 activity response to photoreceptor activation. The reason that HR downregulation and the postsynaptic circuitry response change only started after 3 days remains to be explained. The mechanism underlying the baseline activity of L2 is also unclear. It is broadly believed that the main input that L2 neurons directly receive from photoreceptors is inhibitory because photoreceptors mainly release histamine. However, there is a baseline calcium influx in L2, which seems to be contradictory. In a recently published study, R8 photoreceptors reportedly released not only inhibitory histamine but also excitatory acetylcholine to the postsynaptic neurons and indirectly activated the L2 neurons (Davis et al., 2020). This finding implies that upon light stimulation, L2 is hyperpolarized at first but returns to a slightly depolarized state as the baseline activity after a certain period. This may explain, at least in part, why there is a slight calcium influx in L2 under continuous light conditions even if the main input that L2 neurons directly receive from photoreceptors is inhibitory. Therefore, there may be a threshold of activity for L2, and only after 3 days of LL treatment can calcium influx surpass the threshold and trigger any further changes in the circuit. A previous report also demonstrated that the synapse number in photoreceptors was significantly reduced after constant 3-day light exposure after eclosion in an activity-dependent manner (Sugie et al., 2015). This finding strongly supports the requirement of a 3-day LL condition to pass the activity threshold. It also means that a synapse number decrease in the photoreceptors takes place prior to HR downregulation in the postsynaptic neurons. Upon long-term light exposure, from day 3, the increased calcium influx in L2 triggers HR downregulation. The reduced HR level in the constitutively active L2 neurons attenuates the inhibitory input directly from the photoreceptors and further enhances calcium influx in L2. Therefore, L2 neurons with fewer HRs can respond more drastically to the photoreceptor stimulation after LL treatment.

## Conclusions

In this paper, we report that in the adult *Drosophila* visual system, after constant light exposure-induced synaptic plasticity, inhibitory neurotransmitter histamine receptor transcriptional downregulation occurs in postsynaptic lamina neuron L2, but not in L1. This downregulation depends on the photoreceptor and postsynaptic neuronal activity and involves CaMKII and CREB-B. The histamine receptor transcriptional downregulation results in a more intense postsynaptic L2 neuronal response to subsequent environmental stimulation, especially in the medulla. The results successfully demonstrated the circuit change after synaptic remodeling evoked by long-term activation and demonstrated *in vivo* evidence of the circuitry plasticity upon long-term environmental stimulation. The findings of this research deepen the understanding of the consequences of activity-dependent synaptic plasticity to the circuit and provide new insights into the activity-dependent synaptic plasticity mechanism in the neural circuit.

## Acknowledgements

We gratefully acknowledge Matthew Y. Pecot (Harvard Medical School, USA) for providing the R27G05-FLPG.PEST (attP5) line, and Prof. Chi-Hon Lee (ICOB, Academia Sinica) for UAS-ort lines. We thank Kyoto Stock Center (DGRC), Bloomington *Drosophila* Stock Center, and Vienna *Drosophila* Resource Center (VDRC) for providing fly stocks. We thank Enago (http://www.enago.jp) for the English language review. Y.B. was supported by Japanese Government (MEXT) Scholarship. This work was supported by Grant-in-Aid for Scientific Research on Innovative Areas 16H06457 (T.S.), Grand-in-Aid for Scientific Research (B) 21H02483 (T.S.), Grant-in-Aid for Transformative Research Areas (A) 21H05682 (T.S.) and Takeda Visionary Research Grant from the Takeda Science Foundation (T.S.).

## Reference

Alejevski, F., Saint-Charles, A., Michard-Vanhee, C., Martin, B., Galant, S., Vasiliauskas, D., et al. (2019). The HisCl1 histamine receptor acts in photoreceptors to synchronize Drosophila behavioral rhythms with light-dark cycles. Nat Commun 10(1),252. doi: 10.1038/s41467-018-08116-7.

Bai, Y., and Suzuki, T. (2020). Activity-Dependent Synaptic Plasticity in Drosophila melanogaster. Front Physiol 11, 161. doi: 10.3389/fphys.2020.00161.

Buonarati, O.R., Hammes, E.A., Watson, J.F., Greger, I.H., and Hell, J.W. (2019). Mechanisms of postsynaptic localization of AMPA-type glutamate receptors and their regulation during long-term potentiation. Science Signaling 12(562). doi: ARTN688910.1126/scisignal.aar6889.

Carlezon, W.A., Jr., Duman, R.S., and Nestler, E.J. (2005). The many faces of CREB. Trends Neurosci 28(8),436–445. doi: 10.1016/j.tins.2005.06.005.

Chater, T.E., and Goda, Y. (2014). The role of AMPA receptors in postsynaptic mechanisms of synaptic plasticity. Front Cell Neurosci 8, 401. doi: 10.3389/fncel.2014.00401.

Chen, Y., Akin, O., Nern, A., Tsui, C.Y., Pecot, M.Y., and Zipursky, S.L. (2014). Cell-type-specific labeling of synapses in vivo through synaptic tagging with recombination. Neuron 81(2),280–293. doi: 10.1016/j.neuron.2013.12.021.

Chou, W.H., Hall, K.J., Wilson, D.B., Wideman, C.L., Townson, S.M., Chadwell, L.V., et al. (1996). Identification of a novel Drosophila opsin reveals specific patterning of the R7 and R8 photoreceptor cells. Neuron 17(6),1101–1115. doi: 10.1016/s0896-6273(00)80243-3.

Dau, A., Friederich, U., Dongre, S., Li, X., Bollepalli, M.K., Hardie, R.C., et al. (2016). Evidence for Dynamic Network Regulation of Drosophila Photoreceptor Function from Mutants Lacking the Neurotransmitter Histamine. Front Neural Circuits 10, 19. doi: 10.3389/fncir.2016.00019.

Davis, F.P., Nern, A., Picard, S., Reiser, M.B., Rubin, G.M., Eddy, S.R., et al. (2020). A genetic, genomic, and computational resource for exploring neural circuit function. Elife 9. doi: 10.7554/eLife.50901.

Feinberg, E.H., Vanhoven, M.K., Bendesky, A., Wang, G., Fetter, R.D., Shen, K., et al. (2008). GFP Reconstitution Across Synaptic Partners (GRASP) defines cell contacts and synapses in living nervous systems. Neuron 57(3),353–363. doi: 10.1016/j.neuron.2007.11.030.

Gao, X.J., Riabinina, O., Li, J., Potter, C.J., Clandinin, T.R., and Luo, L. (2015). A transcriptional reporter of intracellular Ca(2+) in Drosophila. Nat Neurosci 18(6),917–925. doi: 10.1038/nn.4016.

Hill, S.J., Ganellin, C.R., Timmerman, H., Schwartz, J.C., Shankley, N.P., Young, J.M., et al. (1997). International Union of Pharmacology. XIII. Classification of histamine receptors. Pharmacol Rev 49(3),253–278.

Ho, V.M., Lee, J.A., and Martin, K.C. (2011). The cell biology of synaptic plasticity. Science 334(6056),623–628. doi: 10.1126/science.1209236.

Kerwin, S.K., Li, J.S.S., Noakes, P.G., Shin, G.J., and Millard, S.S. (2018). Regulated Alternative Splicing of Drosophila Dscam2 Is Necessary for Attaining the Appropriate Number of Photoreceptor Synapses. Genetics 208(2),717–728. doi: 10.1534/genetics.117.300432.

Lee, S.J., Escobedo-Lozoya, Y., Szatmari, E.M., and Yasuda, R. (2009). Activation of CaMKII in single dendritic spines during long-term potentiation. Nature 458(7236),299–304. doi: 10.1038/nature07842.

Lisman, J., Schulman, H., and Cline, H. (2002). The molecular basis of CaMKII function in synaptic and behavioural memory. Nat Rev Neurosci 3(3), 175–190. doi: 10.1038/nrn753.

Luo, J., Shen, W.L., and Montell, C. (2017). TRPA1 mediates sensation of the rate of temperature change in Drosophila larvae. Nat Neurosci 20(1),34–41. doi: 10.1038/nn.4416.

Macpherson, L.J., Zaharieva, E.E., Kearney, P.J., Alpert, M.H., Lin, T.Y., Turan, Z., et al. (2015). Dynamic labelling of neural connections in multiple colours by trans-synaptic fluorescence complementation. Nat Commun 6, 10024. doi: 10.1038/ncomms10024.

Matthews, R.P., Guthrie, C.R., Wailes, L.M., Zhao, X., Means, A.R., and McKnight, G.S. (1994). Calcium/calmodulin-dependent protein kinase types II and IV differentially regulate CREB-dependent gene expression. Mol Cell Biol 14(9),6107–6116. doi: 10.1128/mcb.14.9.6107.

McGuire, S.E., Le, P.T., Osborn, A.J., Matsumoto, K., and Davis, R.L. (2003). Spatiotemporal rescue of memory dysfunction in Drosophila. Science 302(5651),1765–1768. doi: 10.1126/science.1089035.

Okamoto, K., Nagai, T., Miyawaki, A., and Hayashi, Y. (2004). Rapid and persistent modulation of actin dynamics regulates postsynaptic reorganization underlying bidirectional plasticity. Nat Neurosci 7(10),1104–1112. doi: 10.1038/nn1311.

Packard, M., Mathew, D., and Budnik, V. (2003). Wnts and TGF beta in synaptogenesis: old friends signalling at new places. Nat Rev Neurosci 4(2), 113–120. doi: 10.1038/nrn1036.

Raymond, L.A., Blackstone, C.D., and Huganir, R.L. (1993). Phosphorylation of amino acid neurotransmitter receptors in synaptic plasticity. Trends Neurosci 16(4),147–153. doi: 10.1016/0166-2236(93)90123-4.

Satoh, A.K., Xia, H., Yan, L., Liu, C.H., Hardie, R.C., and Ready, D.F. (2010). Arrestin translocation is stoichiometric to rhodopsin isomerization and accelerated by phototransduction in Drosophila photoreceptors. Neuron 67(6),997–1008. doi: 10.1016/j.neuron.2010.08.024.

Song, I., and L. Huganir, R. (2002). Regulation of AMPA receptors during synaptic plasticity. Trends in neurosciences 25(11),578–588.

Sugie, A., Hakeda-Suzuki, S., Suzuki, E., Silies, M., Shimozono, M., Mohl, C., et al. (2015). Molecular Remodeling of the Presynaptic Active Zone of Drosophila Photoreceptors via Activity-Dependent Feedback. Neuron 86(3),711–725. doi: 10.1016/j.neuron.2015.03.046.

Sugie, A., Marchetti, G., and Tavosanis, G. (2018). Structural aspects of plasticity in the nervous system of Drosophila. Neural Dev 13(1), 14. doi: 10.1186/s13064-018-0111-z.

Takemura, S.Y., Bharioke, A., Lu, Z., Nern, A., Vitaladevuni, S., Rivlin, P.K., et al. (2013). A visual motion detection circuit suggested by Drosophila connectomics. Nature 500(7461),175–181. doi: 10.1038/nature12450.

Takemura, S.Y., Karuppudurai, T., Ting, C.Y., Lu, Z., Lee, C.H., and Meinertzhagen, I.A. (2011). Cholinergic circuits integrate neighboring visual signals in a Drosophila motion detection pathway. Curr Biol 21(24),2077–2084. doi: 10.1016/j.cub.2011.10.053.

Talay, M., Richman, E.B., Snell, N.J., Hartmann, G.G., Fisher, J.D., Sorkac, A., et al. (2017). Transsynaptic Mapping of Second-Order Taste Neurons in Flies by trans-Tango. Neuron 96(4), 783–795 e784. doi: 10.1016/j.neuron.2017.10.011.

Wu, X., and McMurray, C.T. (2001). Calmodulin kinase II attenuation of gene transcription by preventing cAMP response element-binding protein (CREB) dimerization and binding of the CREB-binding protein. J Biol Chem 276(3),1735–1741. doi: 10.1074/jbc.M006727200.

Yin, J., Gibbs, M., Long, C., Rosenthal, J., Kim, H.S., Kim, A., et al. (2018). Transcriptional Regulation of Lipophorin Receptors Supports Neuronal Adaptation to Chronic Elevations of Activity. Cell Rep 25(5), 1181–1192 e1184. doi: 10.1016/j.celrep.2018.10.016.

Yuan, Q., Xiang, Y., Yan, Z., Han, C., Jan, L.Y., and Jan, Y.N. (2011). Light-induced structural and functional plasticity in Drosophila larval visual system. Science 333(6048),1458–1462. doi: 10.1126/science.1207121.

